# Three-Dimensional-Mapping of Smooth Muscle Morphogenesis in the Vertebrate Gastrointestinal Tract

**DOI:** 10.1101/2025.07.01.662515

**Authors:** Salomé Ruiz Demoulin, Amandine Falco, Norbert Chauvet, Pascal de Santa Barbara, Sandrine Faure

**Affiliations:** Sandrine Faure, PhyMedExp, University of Montpellier, INSERM, CNRS, 641 avenue Doyen Giraud, 34295 Montpellier, Cedex 5, France

**Keywords:** Gastrointestinal, Smooth muscle, Confocal analyses, Tissue-clearing, 3 Dimension

## Abstract

Gastrointestinal (GI) motility depends on smooth muscle contractions to propel contents through the digestive tract. GI muscle is organized into two perpendicular layers: the inner circular smooth muscle (CSM) and the outer longitudinal smooth muscle (LSM) layers. This organization is important for proper motility. Despite several investigations, GI smooth muscle development remains poorly understood due to several limitations. Previous studies mainly relied on αSMA detection in tissue sections and focused on a single organ or developmental stage. They often overlooked key smooth muscle markers such as CALPONIN and γSMA and largely neglected the LSM layer.

In this study, we combined confocal imaging of whole-mount three-dimensional-cleared embryonic chick guts with heatmap analysis to generate the first spatial atlas of smooth muscle development. We characterize the expression patterns of αSMA, γSMA, and CALPONIN1, thereby mapping the dynamics of the formation and differentiation of CSM and LSM layers along the antero-posterior axis. This study provides a framework for future investigations into the regulatory mechanisms governing smooth muscle patterning and maturation along the GI tract.

## INTRODUCTION

Peristalsis is defined as the coordinated cycles of contraction and relaxation movements that propel luminal contents along the digestive tract. It is essential to the health and well-being of individuals at all ages. Peristalsis is driven by the smooth muscle, whose contractions are regulated by the enteric nervous system (ENS) and the interstitial cells of Cajal (Sanders et al., 2016). Gastrointestinal (GI) smooth muscle is organized into two perpendicular layers: the inner circular smooth muscle (CSM) layer and the outer longitudinal smooth muscle (LSM) layer, with the ENS localized between them (Gabella, 2002; Le Guen et al., 2015). The coordination between both layers, which is established during development, is essential for proper food propulsion, and disruption of this coordination is associated with GI motility disorders (Viti et al., 2023). Despite their importance, the sequential development of smooth muscle layers and their spatial organization have not been thoroughly analyzed.

During development, the primitive gut tube is patterned along the antero-posterior axis into three embryonic domains. This gives rise to anatomically and functionally distinct GI regions, including the stomach, duodenum, small intestine, and colon (Roberts, 2000). The umbilical vessel separates the intestine into pre- and post-umbilical segments. In mammals, the transition between the intestine and the colon is marked by the cecum and by paired ceca in avians.

In the GI tract, smooth muscle cells (SMCs) arise from progenitors derived from the splanchnic mesoderm (McKey et al., 2016). The commitment of mesenchymal cells to the SMC lineage can initially be identified by their elongation and clustering, followed by the early expression of the alpha and gamma isoforms of smooth muscle actin (αSMA and γSMA, respectively), and their later organization into the CSM layer (Gabella, 2002). These cells subsequently undergo differentiation, marked by the expression of CALPONIN1 (a marker of differentiation) (Bourret et al., 2017; Duband et al., 1993; Gimona et al., 1990). A similar two-step mechanism seems to recur to form the LSM layer. This whole differentiation process is conserved across vertebrate species (Huycke et al., 2019; McHugh, 1996; Wallace and Burns, 2005; Yamamoto et al., 1996), and occurs simultaneously with the colonization of the gut by enteric neural crest cells (ENCC), which mainly originate from the vagal neural crest cells (Bourret et al., 2017; Burns and Douarin, 1998; Burns et al., 2000; Le Douarin and Teillet, 1973; Yntema and Hammond, 1954). ENCCs migrate along the gut, from the anterior to the posterior region, proliferate, and play important roles in gut patterning before differentiating into neurons and glial cells of the ENS (Faure et al., 2015; Nagy et al., 2012; Young et al., 2005).

Previous studies, largely based on tissue sections, have looked at when the CSM layer forms; however, these studies have either focused on just one organ (such as the stomach or small intestine) or limited stages of development. The morphogenesis of the LSM remains even less well characterized. Extrapolations from a single muscle layer or specific region to the entire GI tract may lead to inappropriate conclusions. Furthermore, most of previous studies on smooth muscle layer development have only examined αSMA expression while γSMA (encoded by *ACTG2*) is the most abundant and specific smooth muscle actin expressed in the digestive SMCs (Arnoldi et al., 2013). Mutations in the *ACTA2* gene are mainly associated with vascular diseases (Guo et al., 2009), while mutations in *ACTG2* are only reported in GI motility disorders, such as chronic intestinal pseudo-obstruction (CIPO) (Viti et al., 2023).

In the last years, tissue-clearing and three-dimensional (3D) whole-organ imaging approaches have emerged as powerful tools for morphogenesis studies, as they enable high-resolution imaging of organs (Vieites-Prado and Renier, 2021). These techniques have been successfully applied to diverse organs such as the heart, brain, kidney, and lung, but, to our knowledge, not to the GI tract, for the study of smooth muscle.

In this study, we developed whole-mount 3D confocal imaging of cleared embryonic chick guts to generate the first detailed atlas of smooth muscle development. Our findings reveal a process much more complex than expected, with distinct temporal and region-specific patterns. This work represents an important step toward our understanding how smooth muscle forms. Understanding this process is essential for elucidating the origins of GI motility disorders.

## RESULTS AND DISCUSSION

### Spatiotemporal Analysis of CSM Layer Formation in the GI Tract

We first assessed CSM layer formation using αSMA, the common marker used to track the commitment of smooth mesenchymal cells to the SMC lineage. Given its >99% amino acid sequence homology with γSMA (Supplementary Fig. 1A), antibody specificity was validated by Western blotting (Supplementary Fig. 1B).

At early embryonic stages, αSMA was primarily localized in vascular SMCs present in the umbilical vessel located at the junction between the pre- and post-umbilical intestinal segments. The first mesenchymal expression in the GI tract was detected at embryonic day (E) 4.5 (E4.5; Hamburger-Hamilton stages 25), specifically in the post-umbilical intestine (Fig. 1A, panels a and a’). By E5.5 (HH27), αSMA expression intensified, appearing at high level in two distinct regions. The first was a discrete domain at the junction between the proventriculus (glandular stomach) and the gizzard (muscular stomach) (Fig. 1A, panels b and b’). The second covered the intestinal area adjacent to the umbilical vessel, including both pre- and post-umbilical segments (Fig. 1A, panels b and b”). In both regions, staining sharply outlined the developing CSM layer. Lower expression was detected in the colon, indicating a more diffuse and less well-defined CSM layer (Fig. 1A, panels b and b’”). An even more diffuse signal was observed in the gizzard (Fig. 1A, panels b and b’), clearly indicating that the CSM layer had not yet formed in this region at this developmental stage. Western blot analysis of protein extracts from matched gizzard and colonic samples confirmed that αSMA expression begins earlier in the colon than in the gizzard (Supplementary Fig. 2A). Furthermore, the cecum, located between the intestine and the colon, still lacked detectable αSMA expression, indicating a discontinuous pattern of CSM layer emergence beyond the umbilicus (Fig. 1A, panel b).

**Figure 1:**
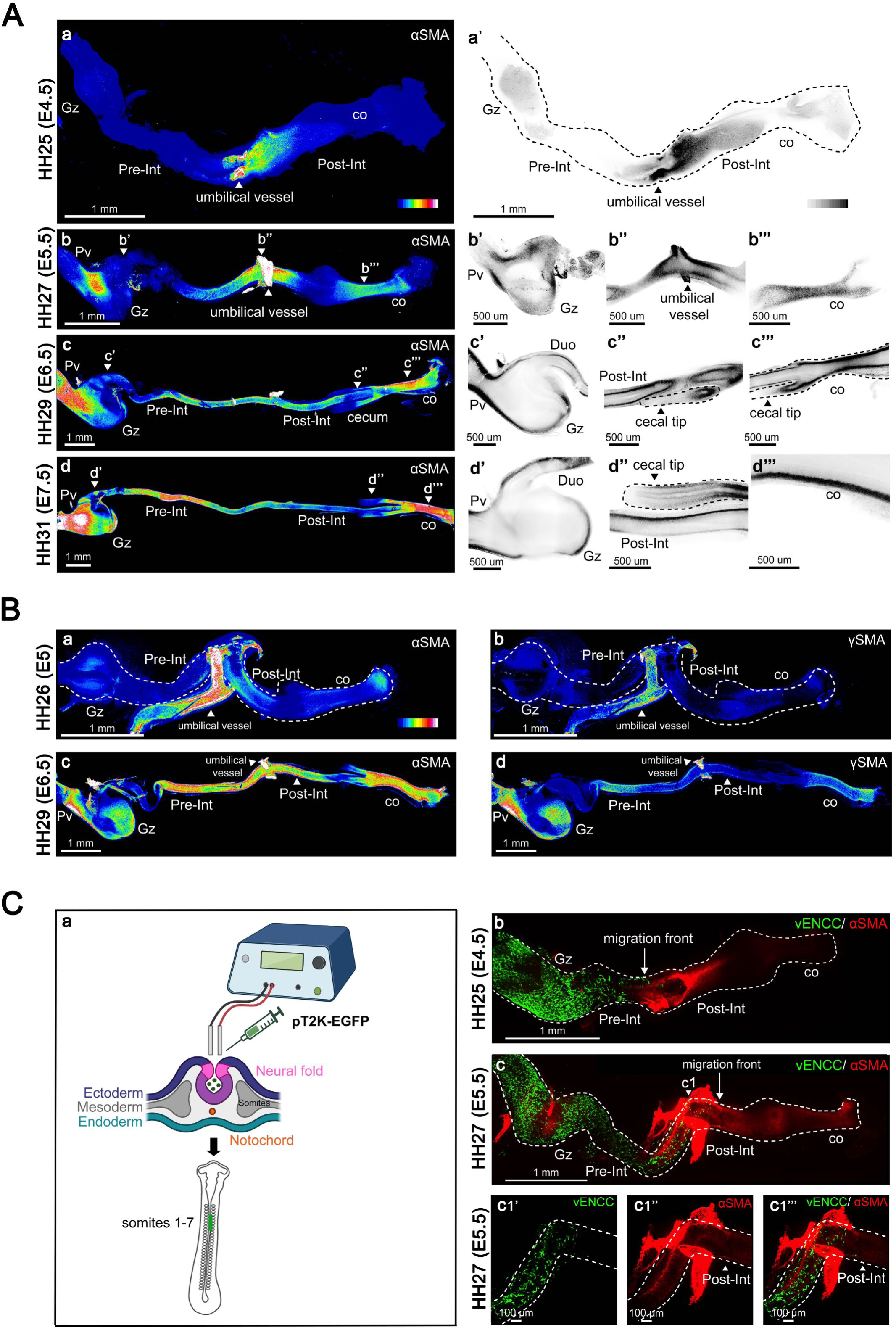
CSM Layer Formation Dynamics in the GI Tract. **(A)** Analysis of αSMA expression. Guts were examined at (a) E4.5 (HH25), (b) E5.5 (HH27), (c) E6.5 (HH29), and (d) E7.5 (HH31). Shown are maximum intensity Z-projection of confocal image stacks of the whole GI tract, represented as heat maps. (a-d) White/red colors indicate high expression levels, while dark blue indicates low expression levels. (a’-d’”) Confocal single Z plane images of the gut regions indicated in panels a-d. Scale bar is indicated for each panel. The dashed lines outline the cecal region in panels c”, c’” and d”. High expression levels are shown in black, and low levels in white. **(B)** Maximum intensity Z-projection of confocal image stacks of guts co-stained for αSMA and γSMA. αSMA staining was observed at (a) E5 (HH26) and (c) E6.5 (HH29); γSMA staining at the same stages is shown respectively in panels (b) and (d). Scale bar is indicated for each panel. **(C)** Analysis of intestinal CSM layer formation in relation to vENCC colonization. (a) Schematic representation of the *in ovo* electroporation procedure to target vagal neural crest cells. (b-c) Maximum intensity Z-projection of confocal image stacks of guts co-stained with anti-GFP and anti-αSMA antibodies at (b) E4.5 (HH25), and (c-c1’”) E5.5 (HH27). Scale bar is indicated for each panel. Abbreviations: Co, colon; vENCC, vagal enteric neural crest cell; Gz, gizzard; Pre-Int, pre-umbilical intestine; Post-Int, post-umbilical intestine; Pv, proventriculus.

By E6.5 (HH29), αSMA expression was mainly observed in the proventriculus, duodenum, pre- and post-umbilical intestine, and in the colon (Fig. 1A, panels c-c’”). While expression was detected in the gizzard, the level remained lower than the signal observed in the colon (Fig. 1A, panel c; see also panels c’ and c’”). Western blots confirmed this difference (Supplementary Fig. 2B). This suggests a less developed CSM layer in the gizzard compared to the colon. αSMA expression was detectable at the base of the cecum from E6.5 (Fig. 1A, panels c, c’’ and c’”) but remained absent from the cecal tips, even at E7.5 (Fig. 1A, panels d and d”). The cecal tips were the last regions of the GI tract to develop the CSM layer, with αSMA appearing there only from E8.5 (HH35) onward (Supplementary Fig. 2C). These data support two main conclusions: first, the CSM layer is initiated around E4.5/E5.5, and is nearly fully established along the GI tract by E6.5, except in the cecal tips, where formation is delayed until E8.5. This delay may reflect a developmental window prioritizing ENS development in the cecum, before their migration into the colon (Nagy et al., 2021). Second, the CSM layer emerges sequentially between E4.5/E5.5 and E6.5, first in the pre- and post-umbilical intestine, then in the proventriculus and colon, and lastly in the gizzard. This finding challenges the previously proposed model that suggests that the commitment of muscle progenitors and, consequently the CSM layer formation, initiates from both anterior and posterior poles of the GI tract (Bourret et al., 2017; Wallace and Burns, 2005).

Next, we analyzed γSMA expression in comparison to αSMA. While αSMA is a well-established marker of mesenchymal progenitor commitment and smooth muscle layer formation, the pattern of γSMA remains less defined. To assess whether γSMA marks progenitor commitment or later stages of differentiation, we performed co-staining experiments. αSMA expression was detectable in the intestinal and colonic mesenchyme at E5, and by E6.5, its staining covered nearly the entire gut (Fig. 1B, panels a and c). In contrast, γSMA was not detected in the gut mesenchyme at E5 and only became detectable from E6.5 onward, primarily in the gizzard, the pre-umbilical intestine, and the colon, exhibiting a discontinuous pattern along the antero-posterior axis (Fig. 1B, panels b and d). Western blot analysis of protein extracts confirmed the onset of γSMA expression at E6.5 (Supplementary Fig. 2D). This demonstrates that γSMA induction is delayed compared to αSMA, indicating that γSMA marks a distinct and later step compared to the commitment step marked by αSMA.

Our data indicate that, instead of forming simultaneously, the CSM layer develops in spatially distinct gut regions, indicating a region-specific regulation of its development. Among the regulatory mechanisms of SMC development are the vENCCs. Indeed, we have previously observed that vENCCs play an important role in smooth muscle commitment in the gizzard (Faure et al., 2015). In chick embryos, vagal neural crest cells emigrate from the neural tube at the level of somites 1-7, enter the stomach by E2.5, reach the intestine-colon junction by E5.5, and the colorectum by E8 (Burns and Douarin, 1998; Le Douarin and Teillet, 1973). We next investigated the onset of αSMA expression in the pre- and post-umbilical intestine relative to the arrival of vENCCs. To do so, we electroporated the pT2K-EGFP plasmid into the neural tube prior to neural crest cell delamination (Fig. 1C, panel a) and analyzed αSMA expression between E4.5 and E5.5. At E4.5, GFP-positive vENCCs, randomly distributed in the mesenchyme of the developing stomach and duodenum, had reached the level of the umbilicus. No αSMA-positive cells were detected in these regions, indicating that the mesenchyme had not yet began to be specified (Fig. 1C, panel b). By E5.5, αSMA-positive cells were detected in the stomach and the pre-umbilical intestine. This suggests that smooth muscle commitment in these regions is initiated after its colonization by vENCCs. In contrast, αSMA was already expressed in the post-umbilical intestine well before the arrival of vENCCs (Fig. 1C, panels c1-c1”’).

### Spatiotemporal Analysis of SMC differentiation in the CSM layer

To determine the timing of SMC differentiation within the CSM layer, we analyzed CALPONIN1 expression and performed co-staining with γSMA. CALPONIN1 was first detected at E6.5 in the gizzard, marking the presence of a well-differentiated CSM layer (Fig. 2A, panels a and a’). Lower levels were observed in the pre-umbilical intestine and the colon (Fig. 2A, panels a, a” and a””). This is surprising as the gizzard is not the first region where the CSM layer forms. Later on, staining becomes more intense along the antero-posterior axis, revealing a clearly differentiated CSM layer in the stomach (proventriculus and gizzard), the pre-umbilical intestine, and the colon from E7 onward (Fig. 2A, panels b and b’). Surprisingly, the CSM layer of the post-umbilical intestine remained undifferentiated at E7 (Fig. 2A, panels b and b’). Signal became detectable at E7.5 (Fig. 2A, panels c-c”). This is surprising, as this region is one of the first where the CSM layer initially formed (Fig. 1A, panel b). Differentiation of the CSM layer in this region becomes clearly evident by E8.5, when CALPONIN1 expression was detected throughout the gut (Fig. 2A, panels d-d’”), including the cecal tips. These observations indicate that differentiation of the CSM layer begins at E6.5 and is completed by E8.5 throughout the GI tract. Strikingly, the timing of SMC differentiation seems uncoupled from the sequential formation of the CSM layer, as regions where the CSM layer forms earliest are not necessarily the first to undergo differentiation. This is particularly evident in the post-umbilical intestine, where the CSM layer forms between E4.5/E5.5, while the differentiation of SMCs is not observed until E8.5. In contrast, in the gizzard, the CSM layer begins to form around E6.5, shortly followed by its differentiation.

**Figure 2:**
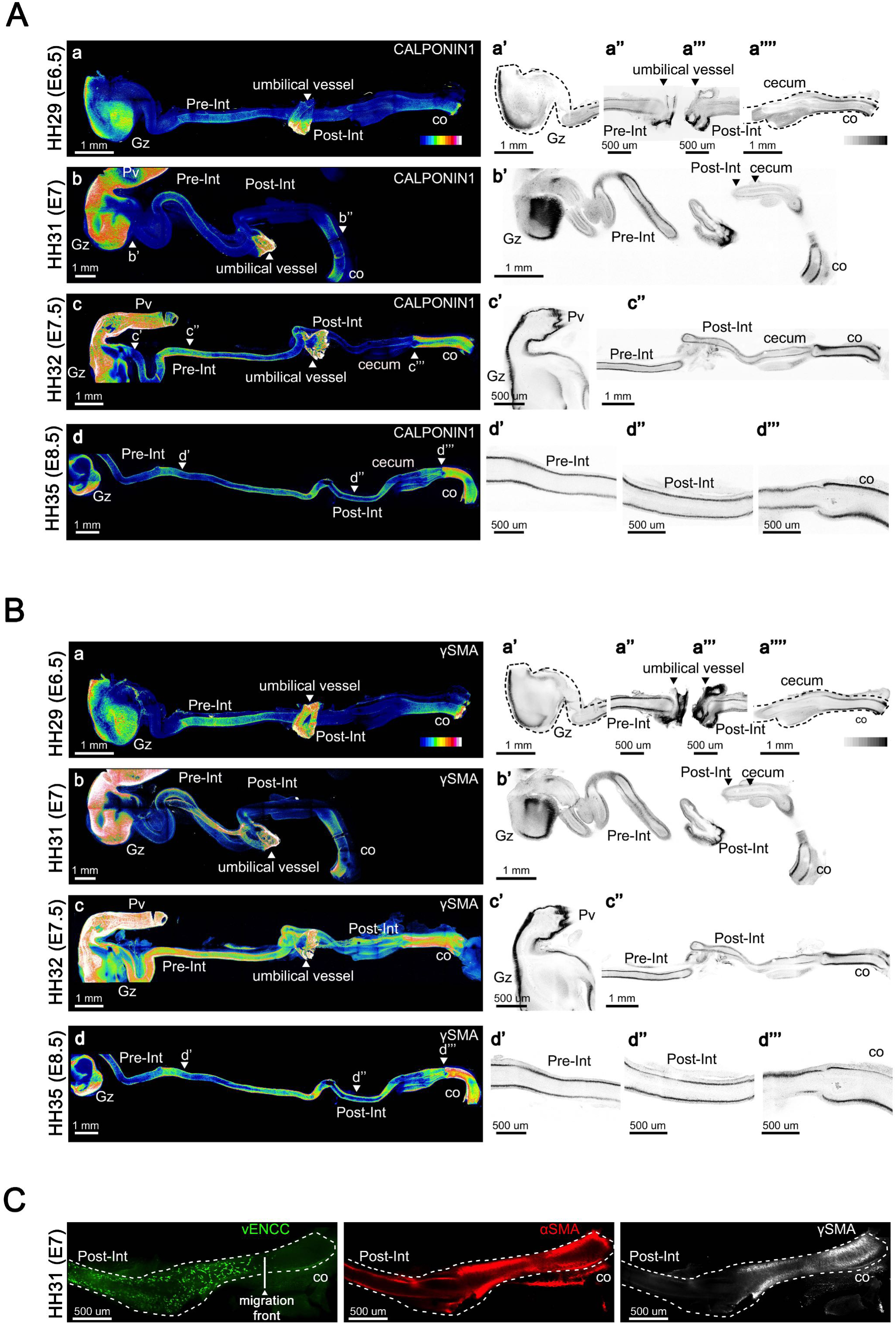
SMC Differentiation Dynamics in the CSM Layer. Guts were co-stained for CALPONIN1 and γSMA. **(A)** Analysis of CALPONIN1 expression. Shown are maximum intensity Z-projection of confocal image stacks of the whole GI tract, represented as heat maps. Guts were examined at (a) E6.5 (HH29), (b) E7 (HH31), (c) E7.5 (HH32), and (d) E8.5 (HH35). (a’-d’’’) Confocal single Z plane images of the level of the gut indicated in panels a-d. Scale bar is indicated for each panel. High expression levels are shown in black, and low levels in white**. (B)** Analysis of γSMA expression on the same guts. (a) E6.5 (HH29), (b) E7 (HH31), (c) E7.5 (HH32), and (d) E8.5 (HH35). (a’-d’’’) Confocal single Z plane images of the level of the gut indicated in panels a-d. Scale bar is indicated for each panel. High expression levels are shown in black, and low levels in white. **(C)** Examination of the intestinal CSM layer differentiation in relation to vENCC colonization at E7 (HH31). Maximum intensity Z-projection of confocal image stacks of guts co-stained with anti-GFP (marking vENCCs), anti-αSMA, and anti-αSMA antibodies. Scale bar is indicated for each panel. Abbreviations: Co, colon; vENCC, enteric neural crest cell; Gz, gizzard; Pre-Int, pre-umbilical intestine; Post-Int, post-umbilical intestine; Pv, proventriculus.

Examination of γSMA expression pattern between E6.5 and E8.5 revealed a profile closer to that of CALPONIN1 than to that of αSMA (compare Fig. 2A and Fig. 2B). γSMA was first observed at E6.5 in the gizzard. Lower levels were observed in the colon and the pre-umbilical intestine (Fig. 2B, panels b and b’). γSMA was not detected in the CSM layer of the post-umbilical intestine at E6.5/E7, and only a faint expression was observed in this region by E7.5 (Fig. 2B, panels c-c’’). As observed for CALPONIN1, γSMA was detected throughout the gut by E8.5 (Fig. 2B, panels d-d’”). The close parallel pattern between γSMA and CALPONIN1 confirms the idea that γSMA marks smooth muscle differentiation.

Although the intestinal CSM layer emerges simultaneously in both pre- and post-umbilical segments, their differentiation timing differs, suggesting region-specific regulation. To investigate this, we analyzed the temporal relationship between CSM layer differentiation and vENCC arrival using lineage tracing (Fig. 2C). At E7, the vENCC migration front had reached the proximal two-thirds of the colon. αSMA is detected in the post-umbilical intestine and the colon. The colonic CSM layer is fully differentiated, as indicated by γSMA expression. In contrast, the CSM layer in the post-umbilical intestine do not express γSMA at this stage. These observations indicate that the CSM layer in the post-umbilical intestine differentiates after the arrival of vENCCs, whereas in the colon, the CSM layer differentiates well before their arrival.

### Spatiotemporal Analysis of CSM and LSM Layer dynamics in the Developing Small Intestine and Colon

We next examined the developmental dynamics of GI smooth muscle layers at later stages. The LSM layer has previously been reported to emerge at E12.5 in the chick intestine (Shyer et al., 2013). We performed co-staining for αSMA/γSMA (Fig. 3) and γSMA/CALPONIN1 (Fig. 4). The orientation of the section planes is illustrated in Figures 3A and 4A. At E12.5, in both the small intestine (pre- and post-umbilical segments) and the colon, the digestive muscle is clearly organized into two layers. In addition to the CSM layer, the LSM layer has formed and is oriented perpendicularly to the CSM layer. In the small intestine, the CSM layer is organized into muscle bundles expressing both αSMA and γSMA (Fig. 3B, panels b1-b1”). Surprisingly, γSMA displays a differential radial distribution within these bundles, being enriched on the abluminal side (Fig. 3B, panels b2’–b2”). In the colon, the CSM layer is also organized into muscle bundles; however, unlike in the small intestine, these bundles express only γSMA (Fig. 3C, panels c1’-c1”). At this developmental stage, the LSM layer is clearly distinguishable in both the small intestine and the colon, but αSMA and γSMA exhibit distinct expression patterns. In the small intestine, the LSM layer is characterized by αSMA expression, as γSMA is not yet detectable (Fig. 3B, panels b2-b2” and b3-b3”). In the colon, in addition to αSMA, a faint γSMA signal was detected in addition to αSMA (compared Fig. 3B and Fig. 3C). This suggests a more advanced stage of differentiation compared to the small intestine. Thus, although the LSM layer is morphologically established by E12.5 in both regions, its molecular differentiation remains incomplete at this stage.

**Figure 3.**
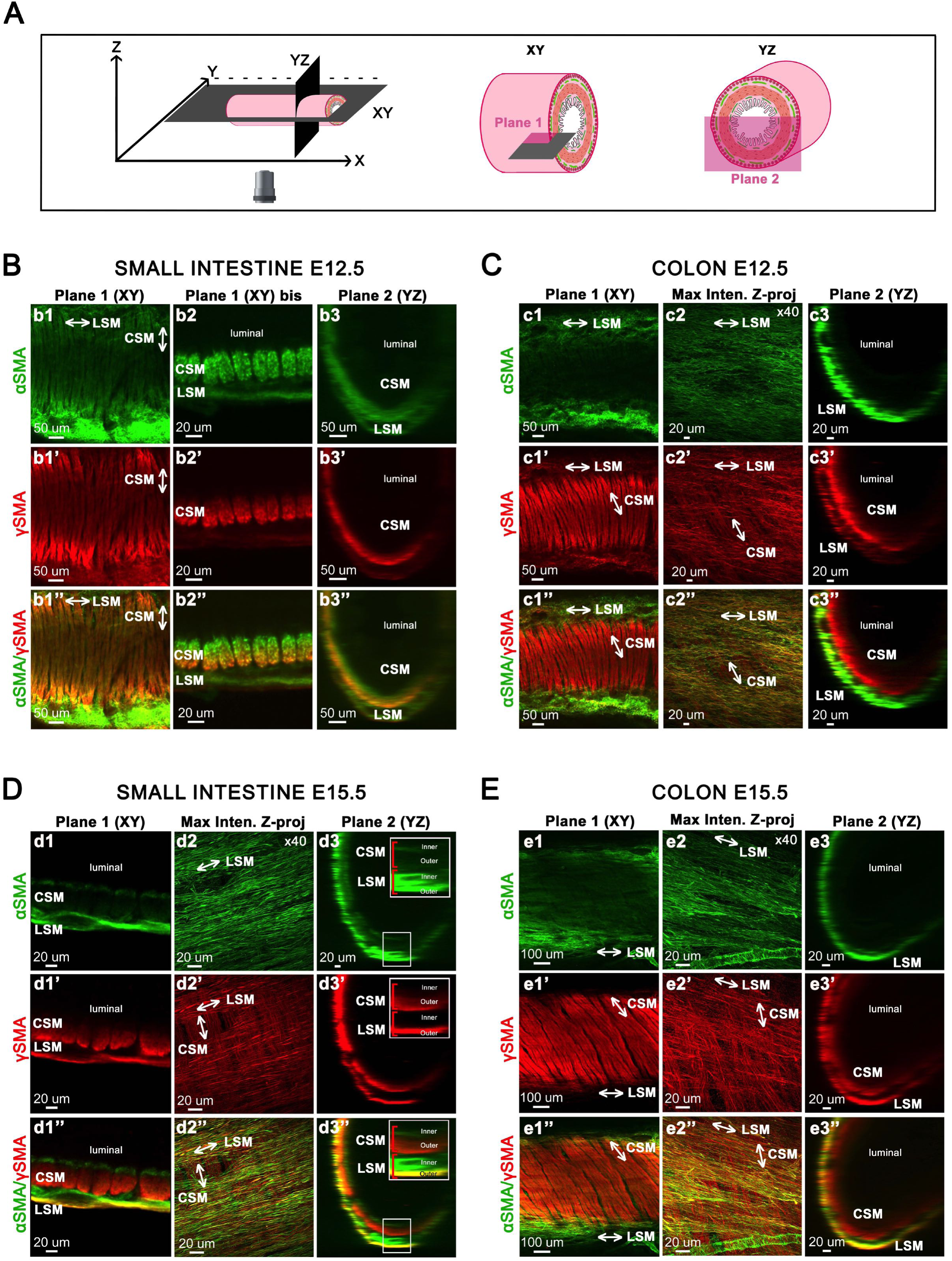
CSM and LSM Layer Dynamics in Developing Small Intestine and Colon at E12.5 and 15.5. **(A)** Schematic representation of digital cross-sectional views of the small intestine and colon. **(B-E)** Guts were co-stained with anti**-**αSMA and anti-γSMA antibodies at E12.5 in the small intestine **(B)** and colon **(C)**, and at E15.5 in the small intestine **(D)** and colon **(E)**. Plane 1 (Confocal single Z plane image) and Plane 2 (transverse section) were obtained from 3D images acquired using an Olympus SR spinning disk confocal microscope with a x10 air objective. Maximum intensity Z-projections were performed using an LSM 800 confocal microscope with a x40 water-immersion objective. Scale bar is indicated for each panel. Are shown in D, panels d3-d3”, focus on different regions of the muscle layers (inner and outer compartments). Scale bar is indicated for each panel. Abbreviations: CSM, Circular Smooth Muscle layer; LSM, Longitudinal Smooth Muscle layer.

**Figure 4.**
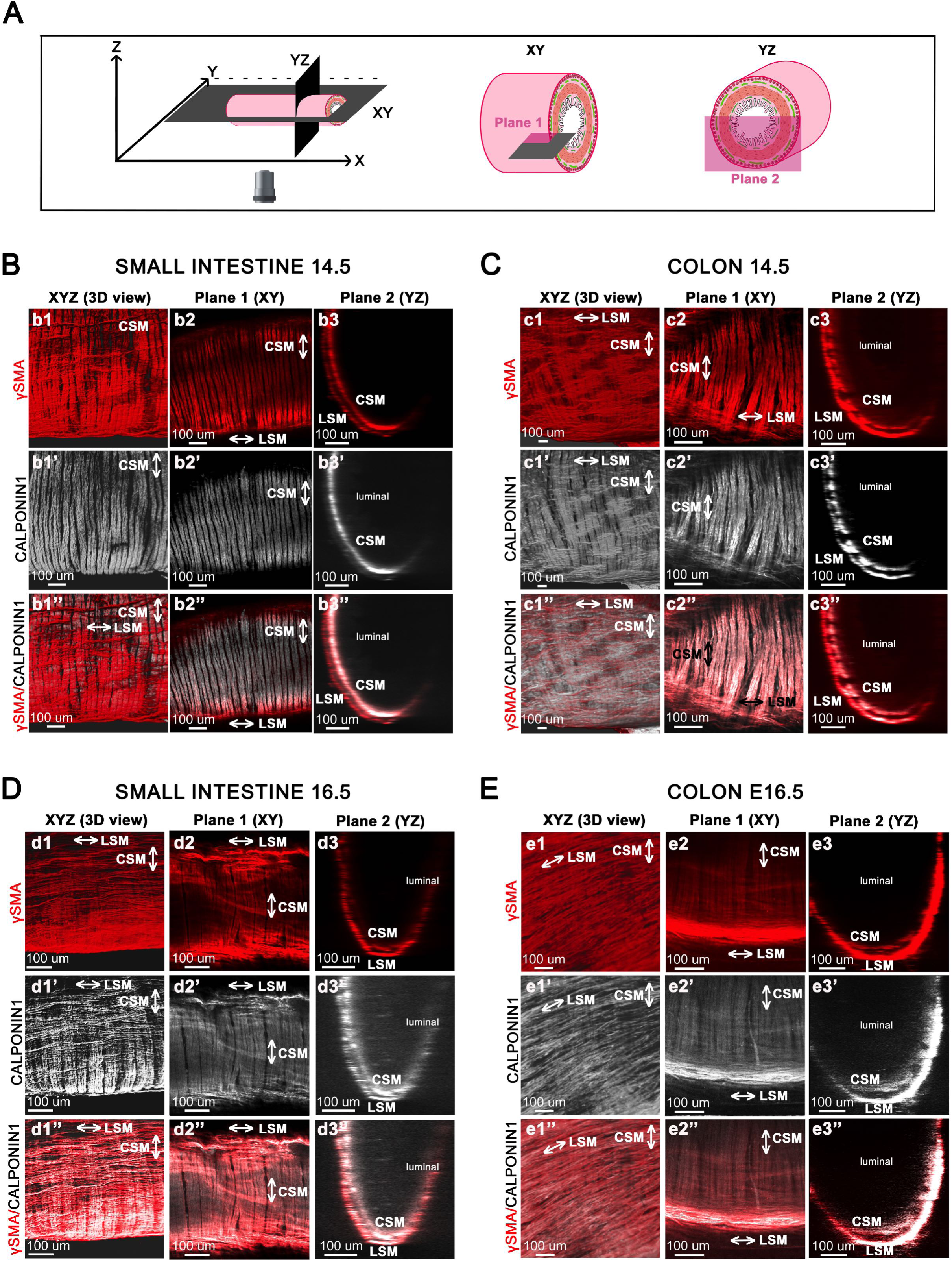
CSM and LSM Layer Differentiation in Developing Small Intestine and Colon at E14.5 and 16.5. **(A)** Schematic representation of digital cross-sectional views of the small intestine and colon. **(B-D)** Guts are co-stained with anti-γSMA and anti-CALPONIN1 antibodies at E14.5 in the small intestine **(B)** and colon **(C)**, and at E16.5 in the small intestine **(D)** and colon **(E)**. Plane 1 (Confocal single Z plane image) and Plane 2 (transverse section) were taken from 3D images obtained with the Olympus SR spinning disk confocal microscope at x10 air objective. Scale bar is indicated for each panel. Abbreviations: CSM, Circular Smooth Muscle layer; Layer; LSM, Longitudinal Smooth Muscle layer.

By E15.5, αSMA expression is almost completely extinguished in the CSM layer of the small intestine. This downregulation was already observed in the colon starting at E12.5 and similarly occurs in the small intestine from E15.5 onward. However, compared to the colon, a very thin line of αSMA expression persists on the luminal side, clearly demarcating the inner compartment of the intestinal CSM layer (Fig. 3D, panels d1 and d3), as previously reported in quail (Thomason et al., 2012). Thus, from this stage, γSMA becomes the predominant smooth muscle-specific actin isoform expressed in the CSM layer of both in the small intestine and colon. In the LSM layer, γSMA is now expressed in the small intestine, similarly to αSMA (Fig. 3D, panels d1’-d3’; panels d1”-d3”). However, αSMA and γSMA display distinct expression patterns: αSMA is present in both the inner and outer compartments, whereas γSMA is restricted to the outer compartment (Fig. 3D, panels d1”-d3”). In the colon, both layers are well developed and γSMA is expressed in both layers, while αSMA expression is confined to the LSM layer (Fig. 3E).

Our data suggest that the LSM of the colon differentiates earlier than that of the small intestine. Analysis of CALPONIN1 expression further supports this regional difference, as it emerges earlier in the colonic LSM at E14.5 (compare Fig. 4B, panels b1’-b3’ to Fig. 4C, panels c1’-c3’). CALPONIN1 was not detected in the intestinal LSM layer at that stage, but became evident at E16.5 (Fig. 4D panels d1’-d3’).

## MATERIALS AND METHODS

### Animals

Brown chicken eggs that had been fertilized (Les Bruyères, Dangers, France) were kept in a humidified incubator (Ducatillon or Coudelou, France) at 37.3°C. Experiments on embryos were conducted in accordance with institutional ethical guidelines established by INSERM and CNRS. Under EU Directive 2010/63/EU, embryos are exempt from ethical approval. Embryos were staged using the Hamburger and Hamilton stages (Hamburger and Hamilton, 1951). GI tissues were dissected at defined embryonic days (E) (Southwell, 2006), fixed overnight at 4°C in 4% paraformaldehyde, washed in PBS for 1 hour at room temperature, and stored at 4°C prior to whole-mount immunostaining.

### Antibodies

The following primary antibodies were used: mouse anti-αSMA (clone 1A4, Santa Cruz, Cat#sc-32251, RRID: AB_262054), rabbit anti-γSMA (MyBioSource, Cat#MBS820899, RRID: AB_3697683), mouse anti-CALPONIN1 (OriGene Cat#AM20594PU-N, RRID: AB_3697684), goat anti-GFP (Rockland, Cat#600-101-215, RRID: AB_11181883), rabbit MYC (Sigma-Aldrich, Cat#C3956, RRID: AB_439680) and rabbit GAPDH (Sigma-Aldrich, Cat#G9545, RRID: AB_796208).

### Western Blotting

GI tissues were lysed in buffer (20 mM Tris pH 8, 50 mM NaCl, 1% NP40, plus cOmplete™ EDTA-free Protease Inhibitor Cocktail, Roche). Total protein lysates (10 μg) were boiled in SDS sample buffer, separated by 10% SDS-PAGE, and transferred onto nitrocellulose membranes. Blots were probed with either mouse anti-αSMA or rabbit anti-γSMA (all at 1:1000 dilution). Detection was performed using infrared-labeled secondary antibodies and the Odyssey® imaging system (LI-COR). Revert™ Total Protein Stain was used for normalization. Band intensities were quantified using Image Studio™ software (LI-COR), with total protein signal serving as a loading control.

### Immunofluorescence, Tissue Clearing, and Confocal Microscopy

GI tissues were permeabilized for two days at room temperature (RT) in 2% Triton X-100 in PBS (with 0.05% sodium azide). After that, they were washed three times for 15 minutes each time in PBS. Blocking was carried out for two days at 4°C in PBS that contained 0.05% sodium azide, 1% Triton X-100, and 10% donkey serum. Primary antibodies incubated on tissues for three days at 4°C (all at 1:300 dilution). Samples were washed in PBS with 3% NaCl and 0.2% Triton X-100 for 1.5 days at 4°C following primary incubation. Alexa Fluor-conjugated secondary antibodies (anti-rabbit 647 and anti-mouse 568, Invitrogen, 1:300) were incubated for two days at 4°C. Final washes were performed in PBS with 3% NaCl and 0.2% Triton X-100 and incubating overnight with RapiClear® at RT. Samples were mounted in 2,2′-thiodiethanol (Sigma Aldrich #166782) and imaged using a spinning disk (Olympus SR, 10X) or conventional confocal microscope (LSM800, 40X). Images were processed using Fiji and Imaris software.

### In Ovo Electroporation

Fertilized eggs were incubated until stage HH9. Plasmids pCAGGS-T2TP and pT2K-CAGGS-EGFP (Wang et al., 2011) were microinjected into the neural tube at somites 1-7 to allow stable expression of the GFP. Deposit of plasmids in the neural tube was visualized with Phenol Red. Bilateral electroporation was performed using gold-plated electrodes (ECM 830, BTX), delivering three 5 ms pulses at 15 V. Eggs were sealed and re-incubated at 37.3°C. Colons exhibiting GFP expression were selected for subsequent immunostaining.

## Abbreviations used in this paper

3D: three dimensional
αSMA: alpha smooth muscle actin
Co: colon
CSM: Circular Smooth Muscle
E: embryonic day
ENS: enteric nervous system
ENCC: enteric neural crest cell
γSMA: gamma smooth muscle actin
GI: gastrointestinal
Gz: gizzard
LSM: Longitudinal Smooth Muscle
Post-Int: post-umbilical intestine
Pre-Int: pre-umbilical intestine
SMC: smooth muscle cell
vENCC: vagal enteric neural crest cell

## Author Contributions

Conceptualization: S.F., P.d.S.B; Methodology: S.R.D., A.F., N.C.; Formal analysis: S.R.D, S.F.; Investigation: S.F., P.d.S.B. S.R.D; Writing - original draft: S.F., S.R.D; Writing - review & editing: S.F., P.d.S.B., S.R.D., A.F., N.C.; Visualization: S.R.D., A.F.; Supervision: S.F.; Project administration: S.F; Funding acquisition: S.F., P.d.S.B.

## Acknowledgments

The authors thank the members of the “Development of Visceral Smooth Muscle and Associated Pathologies” team (PhyMedExp) for their critical review of the manuscript. They also acknowledge O. Faklaris and V. Becker from the Montpellier Ressources Imagerie (MRI) facility (Biocampus, Montpellier) for providing access to confocal microscopy and technical training.

## Legends of supplementary Figures

**Figure 1.**
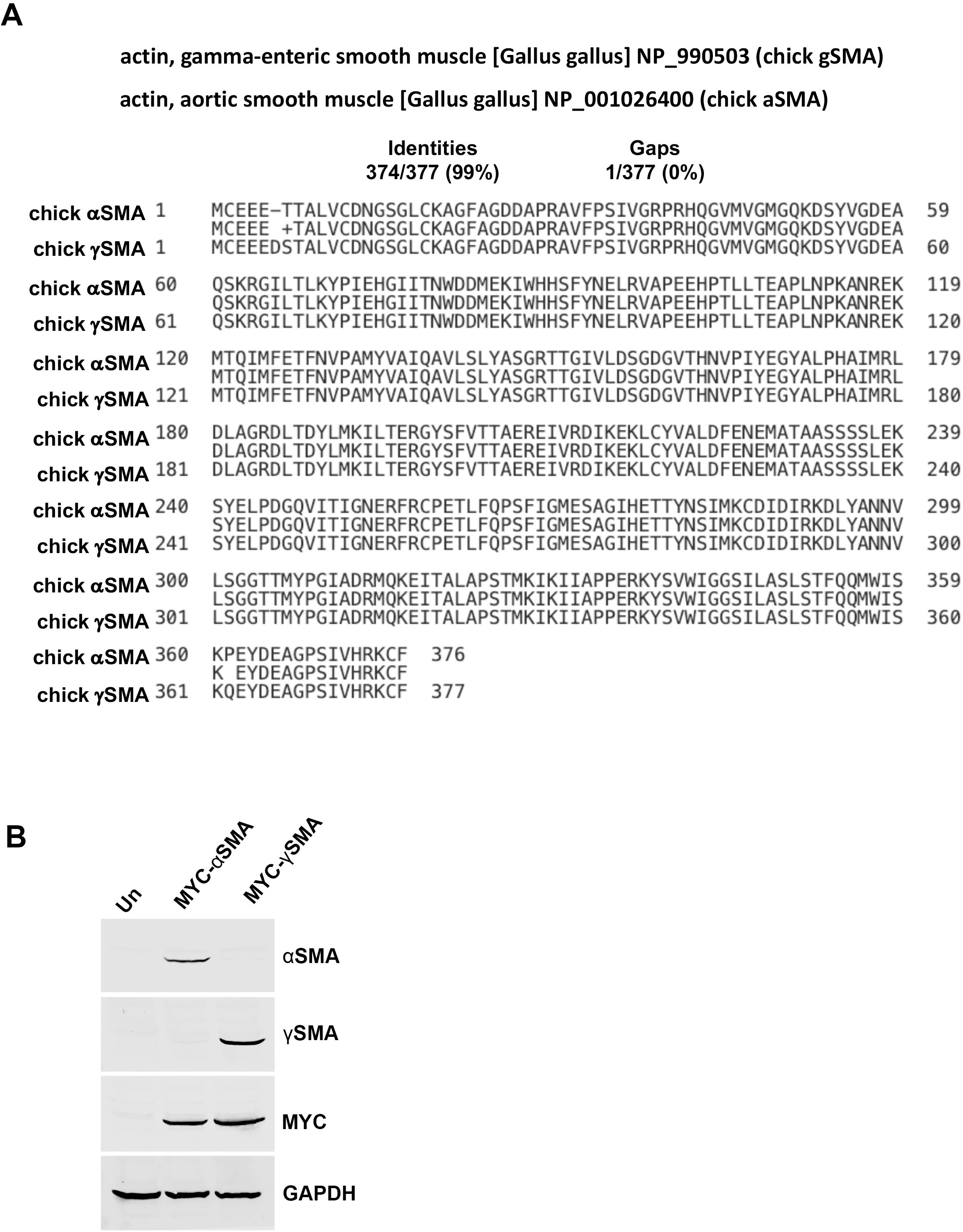
Validation of the specificity of αSMA and γSMA antibodies. **(A)** Alignment of the amino acid sequences of chick αSMA and γSMA proteins. **(B)** Western blot analysis of extracts from HEK293 cell transiently transfected with constructs encoding human αSMA or γSMA, each fused to a MYC tag (MYC-αSMA and MYC-γSMA). Blots were probed with anti-αSMA, anti-γSMA, GAPDH and anti-MYC antibodies. GAPDH level serves as a loading control while MYC level serves as a control for transfection efficiency.

**Figure 2.**
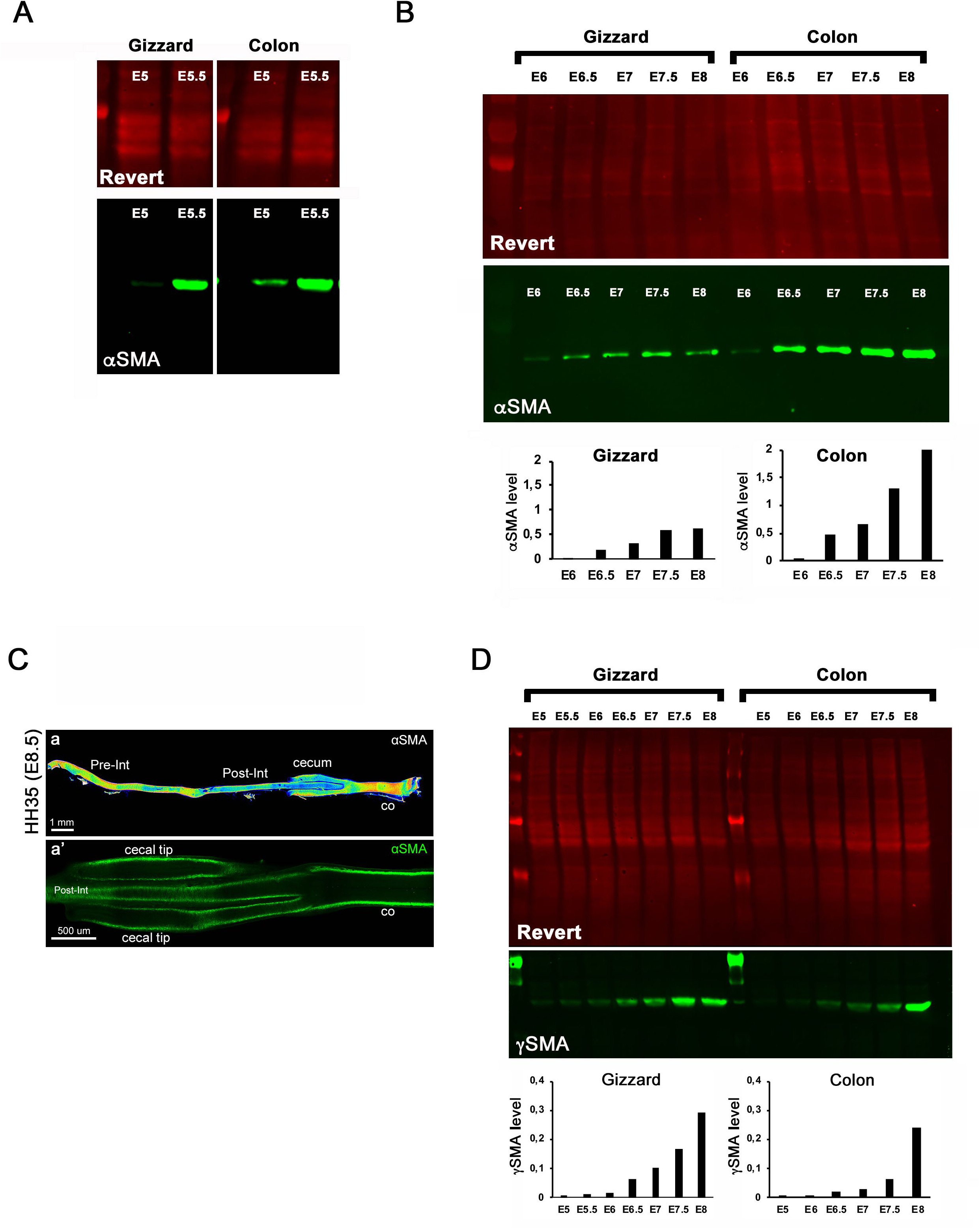
Analysis of αSMA and γSMA expression. **(A)** Representative western blot of gizzard and colonic extracts at E5 and E5.5 probed for αSMA and Revert to evaluate total protein level. **(B)** Representative western blot analysis of gizzard and colonic extracts collected between E6 and E8, probed for αSMA. αSMA levels were normalized to Revert total protein staining. **(C)** Analysis of αSMA expression in chick GI tract at E8.5. (a) Maximum intensity Z-projections of confocal image stacks of the entire GI tract shown as heat maps; (a’) representative single Z-plane confocal image. Abbreviations: Pre-Int, pre-umbilical intestine; Post-Int, post-umbilical intestine; co, colon. **(D)** Representative western blot showing γSMA levels in samples analyzed in (B). γSMA levels were normalized to Revert total protein staining.

